# Increased excursions to functional networks in schizophrenia in the absence of task

**DOI:** 10.1101/2021.11.25.469834

**Authors:** Miguel Farinha, Conceição Amado, Joana Cabral

**Affiliations:** Instituto Superior Técnico, University of Lisbon, Lisbon, Portugal; Department of Mathematics and CEMAT, Instituto Superior Técnico, University of Lisbon, Lisbon, Portugal; Life and Health Sciences Research Institute, School of Medicine, University of Minho, Braga, Portugal; Center for Eudaimonia and Human Flourishing, Linacre College, University of Oxford, Oxford, United Kingdom; Center for Music in the Brain, Department of Clinical Medicine, Aarhus University, Denmark

**Keywords:** Resting-state functional magnetic resonance imaging, Dynamic functional connectivity, LEiDA, Functional networks, Dynamical Systems Theory, Schizophrenia

## Abstract

Brain activity during rest has been demonstrated to evolve through a repertoire of functional connectivity (FC) patterns, whose alterations may provide biomarkers of schizophrenia - a psychotic disorder characterized by dysfunctional brain connectivity. In this study, differences between the dynamic exploration of resting-state networks using functional magnetic resonance imaging (fMRI) data from 71 schizophrenia patients and 74 healthy controls were investigated using a method focusing on the dominant fMRI signal phase coherence pattern at each time point. Through the lens of dynamical systems theory, brain activity in the form of temporal FC state trajectories was examined for intergroup differences by calculating the fractional occupancy, dwell time, limiting probability of each state and the transition probabilities between states. Results showed reduced fractional occupancy of a globally synchronized state in schizophrenia. Conversely, FC states overlapping with canonical functional subsystems exhibited increased fractional occupancy and limiting probability in schizophrenia. Furthermore, state-to-state transition probabilities were altered in schizophrenia. This revealed a reduced probability of remaining in a global integrative state, increased probability of switching from this state to functionally meaningful networks and reduced probability of remaining in a state related to the Default Mode network. These results revealed medium to large effect sizes. Finally, this study showed that using *K*-medoids clustering did not influence the observed intergroup differences - highlighting the utility of dynamical systems theory to better understand brain activity. Combined, these findings expose pronounced differences between schizophrenia patients and healthy controls - supporting and extending current knowledge regarding disrupted brain dynamics in schizophrenia.

## Introduction

Functional magnetic resonance imaging (fMRI) data can be used to characterize brain activity at “rest” as the time-resolved emergence and dissolution of functionally meaningful networks, in other words, the constant reconfiguration of resting-state functional connectivity (FC) patterns or states over time (1). These resting-state networks (RSNs) have been extensively analyzed across neuroimaging studies (2) and their characterization in the temporal domain has been suggested to provide potential biomarkers of several disorders (3–5). Schizophrenia (SZ) is a chronic brain disorder typified by disruption to thought processes, perception, cognition and behaviours, for which there is still a lack of biomarkers. To date, previous studies investigating dynamic functional connectivity (dFC) have suggested that compared to healthy controls (HCs), patients with SZ spend more time in FC states characterized by weak connectivity (6) and less time in FC states which represent strong, large-scale brain connectivity (7–9). Furthermore, when SZ patients transition into the FC state of strongest connectivity, they switch states very rapidly (6). Overall, SZ patients have been found to exhibit fewer changes between connectivity patterns compared to HCs (10).

Most research on dFC in SZ has been carried out using independent component analysis to extract time courses of networks which were subsequently used to estimate dFC through sliding-window analysis (SWA) (3, 5). However, the choice of the window length affects the temporal resolution of the SWA approach - raising questions over its validity (5). In this study, to overcome this weakness, the Leading Eigen-vector Dynamics Analysis (LEiDA) method, based on Blood Oxygenation Level Dependent (BOLD) phase coherence, is used to investigate dFC at an instantaneous level (11, 12). There are three primary aims of this study: (1) assess whether intergroup differences exist by estimating and characterizing time courses of recurrent FC states using tools from dynamical systems theory; (2) examine the validity of the partitions resulting from the clustering procedure; (3) understand the influence of using the *K*-medoids algorithm instead of the *K*-means algorithm on the ability to differentiate SZ patients from HCs. This work hypothesized to find abnormal dFC in SZ patients characterized by: (1) reduced excursions to a FC state possibly involved in the integration of segregated functional connections; (2) increased excursions to a number of FC states which represent functionally segregated networks.

## Materials & Methods

### Neuroimaging data

Neuroimaging data was obtained from the publicly available repository COBRE preprocessed with NIAK 0.17 - lightweight release (13, 14). The neuroimaging data included preprocessed resting-state fMRI (rs-fMRI) data from 72 SZ patients and 74 HCs. The rs-fMRI data featured 150 echo planar imaging BOLD volumes obtained in 5 minutes, with repetition time (TR) = 2 s, echo time = 29 ms, acquisition matrix = 64 × 64 mm^2^, flip angle = 75° and voxel size = 3×3×4 mm^3^.

Inspection of the BOLD data for each subject resulted in the exclusion of one subject whose data did not include all 150 BOLD volumes. Therefore, the final dataset used in this analysis included 71 SZ patients (80.28% males) and 74 HCs (68.92% males). Both groups had an age range of 18-65 years old. A two-sided Wilcoxon Rank-Sum test with Bonferroni correction did not identify a significant difference between the mean age of the groups (*p* = 0.4253). The framewise displacement (FD) provided a quantitative indication of each subject’s head motion during the scanning period (15). The same statistical test detected a significant intergroup difference in the group mean FD (*p <* 0.001). Specifically, on average, the BOLD signals of SZ patients were characterized by larger amounts of head motion (FD).

The acquisition and preprocessing of the data are fully described in detail in (14). Here, contrary to the typical rs-fMRI preprocessing procedure (16), the neuroimaging data was not subject to spatial smoothing, temporal filtering and nuisance regression.

### Parcellation

Following the methodology used by (12, 17– 19), the canonical Anatomic Automatic Labeling (AAL) template was used to parcellate the entire brain of each participant into 90 cortical and sub-cortical non-cerebellar regions. Accordingly, for each region in the brain, at each time point (TR), the BOLD signals were averaged over all voxels belonging to that brain area to compute the regional BOLD time courses. For each subject, this resulted in a *N*×*T* BOLD dataset, where *N* = 90 is the number of brain areas and *T* = 150 is the number of volumes in each scan.

### Computation of dynamic functional connectivity

To compute the phase relationship between each pair of AAL regions, first the instantaneous phase of the BOLD signals across all brain regions *n* ∈ {1, …, *N*} for each time *t* ∈ {2, …, *T*− 1}, *θ*(*n, t*), were estimated by computing the Hilbert transform of their BOLD regional time courses (11). Here, the first and last TR of each fMRI scan were excluded due to possible signal distortions induced by the Hilbert transform (19). The Hilbert transform enables the capture of the time-varying phase of a BOLD signal at each time, *t*, by converting it into its analytical representation (see Figure 1A (top left)) (11, 12). To obtain a whole-brain pattern of BOLD phase synchrony, the phase coherence between areas *n* and *p* at each time *t, dFC*(*n, p, t*), was estimated using Eq. (1):

**Figure 1.**
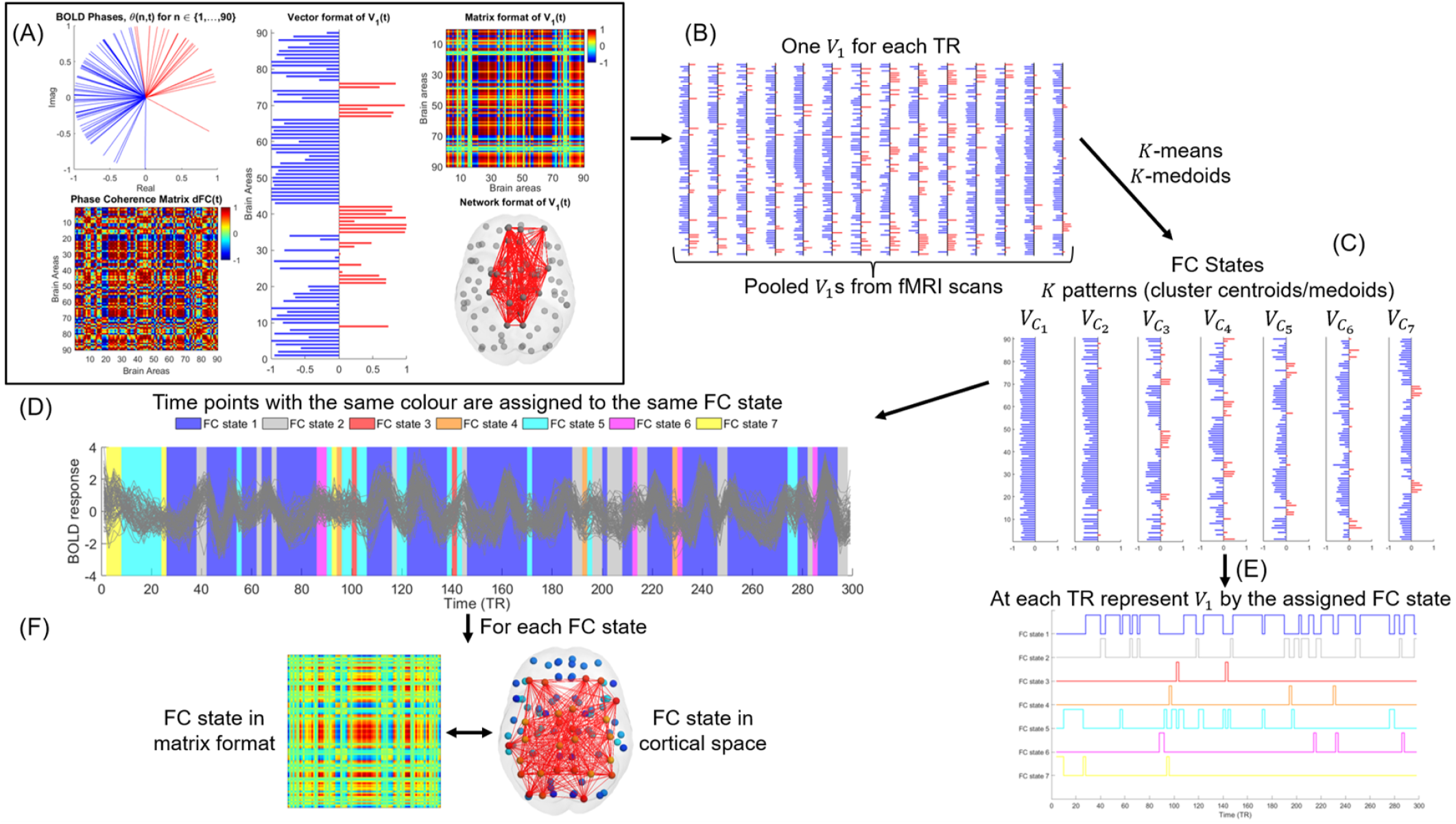
Graphical illustration of the estimation and characterization of the temporal trajectories of recurrent FC states obtained by using Leading Eigenvector Dynamics Analysis (LEiDA). **(A)** BOLD phases of all *N* = 90 brain areas in the complex plane at time *t* (top left); BOLD phase coherence matrix at time *t, dF C*(*t*) (bottom left); Vector representation of the leading eigenvector, *V*_1_ (*t*), of *dF C*(*t*) (middle); Matrix representation of *V*_1_ (*t*) (top right); Network representation of *V*_1_ (*t*), with links between the areas with positive elements in *V*_1_ (*t*) plotted in red (bottom right). **(B)** The leading eigenvectors are computed for each time point and from all fMRI scans. **(C)** The pooled leading eigenvectors are partitioned into *K* clusters using a clustering algorithm. The cluster centroids/medoids are assumed to represent recurrent patterns of BOLD phase coherence (FC states). **(D**,**E)** The leading eigenvector at each TR is represented by the centroid/medoid of the cluster to which it was assigned by the clustering procedure. This originates time courses of FC states for each fMRI session. The time courses are then characterized using tools from dynamical systems theory. **(F)** Each FC state can be represented as a *N* × *N* matrix (outer product) and as a network in cortical space (elements with positive sign linked by red edges).

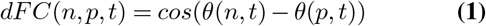

where phase coherence values range between -1 (areas *n* and *p* in anti-phase at time *t*) and 1 (areas *n* and *p* have synchronized BOLD signals at time *t*), as shown in Figure 1A (bottom left). This computation was repeated for all pairwise combinations of brain areas (*n, p*), with *n, p* ∈ {1, …, 90}, at each time point *t*, with *t* ∈ {2, …, 149}, and for all subjects. For each subject, the resulting dFC was a three-dimensional tensor with dimension *N* × *N* × *T* ^′^, where *T* ^′^ = 148, i.e., 148 *dFC*_90×90_(*t*) matrices were estimated.

### Functional connectivity leading eigenvector

To characterize the evolution of the phase coherence matrix over time with reduced dimensionality, the current study employed the LEiDA method which considers only the leading eigenvector, *V*_1_(*t*), of each *dFC*(*t*) matrix (12). In detail, as observed in Figure 1A (middle), the leading eigenvector, *V*_1_(*t*), is a *N*×1 vector that captures the dominant connectivity pattern of BOLD phase coherence at time *t*, i.e., *V*_1_(*t*) represents the main orientation of the BOLD phases over all brain areas (12). Under this framework, for each time *t*, the associated leading eigenvector partitions the *N* brain areas into two communities by separating the elements with different signs in *V*_1_(*t*) (12, 20). When all elements of *V*_1_(*t*) have the same sign, the BOLD phases between brain regions are coherent, which is indicative of a global mode of phase coherence governing all BOLD signals. This implies that all brain regions belong to the same community. Contrarily, if the elements of *V*_1_(*t*) have different signs (i.e., positive and negative), the connectivity pattern between brain regions is not coherent. As a result, each brain area is assigned to one of the two communities according to their BOLD phase relationship. Additionally, the absolute value of each element in the leading eigenvector weighs the contribution of each brain area to the assigned community (12, 20). The dominant FC pattern of the *dFC* matrix at time *t* can also be reconstructed back into matrix format by computing the (*N*×*N*) outer product 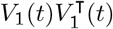, as shown in Figure 1A (top right). Given that if *V*_1_(*t*) is a leading eigenvector, so is − *V*_1_(*t*), following the procedure of (17–19), it was ensured that most of the elements in *V*_1_(*t*) had negative values. This is because by assigning positive values to the brain areas whose BOLD phases did not follow the global mode, functional brain networks were distinctly detected, as seen in Figure 1A (bottom right). Importantly, this approach was found to explain most of the variance of observed BOLD phase coherence data variation, while substantially reducing its dimensionality. In fact, the leading eigenvector accounted for more than 50% of the variance in phase coherence at all time points and for all subjects.

### Estimation of FC states

Upon computing the leading eigenvector of the phase coherence matrix for each recording frame, the next step in the analysis was to characterize the evolution of the dFC over time by identifying recurrent FC states in the data, as illustrated in Figure 1C (12).

Firstly, the conventional LEiDA clustering analysis was performed by applying the *K*-means algorithm (21) to the dataset of all leading eigenvectors computed at the set of volumes {2, …, 149} across all 145 participants (see Figure 1B). This corresponded to a combined total of 148 × 145 = 21460 leading eigenvectors. Lastly, the impact of using the *K*-medoids algorithm (21) to conduct a LEiDA analysis was investigated through its application to the combined total of 21460 leading eigenvectors. Here, both algorithms were run with a value of *K* from 2 to 20, i.e., dividing the set of leading eigenvectors into *K* = {2, 3, …, 20} clusters. Furthermore, in both clustering analyses, the cosine distance was used as the distance metric for minimization and the algorithms were run 1000 times to minimize the chances of getting trapped in a local minima (12, 17–19).

### Characterization of FC state trajectories

Independently of the algorithm, the LEiDA clustering procedure outputs one distinct clustering solution for each value of *K* clusters. Specifically, each clustering solution contains *K* clusters **C** = {*C*_1_, …, *C*_*K*_}, with *K* ∈ {2, …, 20}- decomposing the *N* -dimensional phase space of pooled leading eigenvectors into a *K*-dimensional state space. Let *V*_*C*_*α* be the vector of dimension *N*×1 representing the centroid/medoid of cluster *C*_*α*_, where *α* ∈ {1, …, *K*}. Then, for a given clustering solution, each *V*_*C*_*α* represents a recurrent FC state, as depicted in Figure 1C (12).

For each clustering solution, the set of estimated *K* FC states was used to obtain, for each participant, time courses of FC states (as represented in Figure 1D,E). This was accomplished by representing each *V*_1_ at time *t* by the FC state (centroid/medoid) of the cluster to which it was assigned by the clustering algorithm. Specifically, following the conceptual framework proposed by (19), resting-state BOLD time series were assumed to temporally evolve through a finite state trajectory of recurrent patterns of BOLD phase co-herence. Following this rationale, each clustering solution with *K* FC states was assumed to define a finite state space *S* = {1, …, *K*}. Furthermore, for a clustering solution with *K* clusters, the cluster (FC state) to which *V*_1_(*t*) was assigned at time *t*, denoted by *V*_*t*_, was assumed to define a stochastic process, {*V*_*t*_ : *t* ∈ {2, …, 149}}, with an associated finite state space given by *S*. Consequently, considering the Markov property (22) holds, each temporal trajectory of FC states was assumed to define a time-homogeneous Discrete Time Markov Chain (DTMC). Importantly, it must be noted that, although brain activity is an uninterrupted process, the restricted fMRI scanning windows implied the state trajectories were temporally limited - resulting in a number of DTMCs not spanning the entire state space.

A number of descriptive measures were considered to characterize the properties of the temporal trajectories of FC states observed in SZ patients and HCs. Notably, these measures have been shown to provide relevant insights on aberrant dynamic brain activity in previous LEiDA analyses (12, 17, 18).

#### Fractional occupancy

The fractional occupancy (probability of occurrence) of a FC state *α* represents the proportion of times *V*_*t*_ is assigned to cluster *C*_*α*_ throughout a scan (19). The fractional occupancy of FC state *α* for the fMRI scan of subject 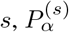, is calculated (estimation) as follows:

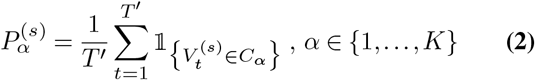

where *T* ^′^ = 148 is the number of time points (first and last volume of each scan were excluded), 𝟙 is the indicator function and 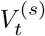 is the FC state to which *V*_1_(*t*) was assigned at time *t*. For each clustering solution, this measure was estimated for each of the *K* FC states separately for each fMRI scan.

#### Dwell time

The dwell time (mean duration) of a FC state represents the mean number of consecutive epochs spent in that state throughout the duration of a scan (19). The dwell time of FC state *²*,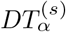, is defined (estimation) as:

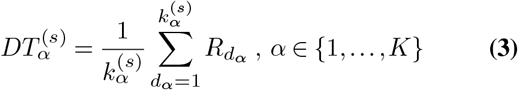

Where 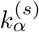 is the number of consecutive periods in which 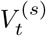 was assigned to cluster *C*_*α*_ and 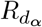 is the duration of each of the 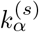 periods. For each clustering solution, the dwell time was estimated for each of the *K* FC states separately for each fMRI scan.

#### One-step transition probability matrix

According to (19), considering a clustering solution with state space *S* = {1, …, *K*}, the probability of being in FC state *α* at time *t* and transition to FC state *β* at time *t* + 1 is given by the following expression:

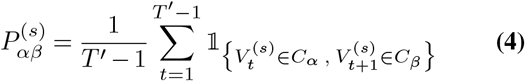

with *α*, ∈ *β* {1, …, *K*}. From Eq. (4), for a clustering solution with *K* FC states, it follows that the Transition Probability Matrix (TPM) of the fMRI scan of subject *s*, **P**^(*s*)^, is defined (estimation) as:

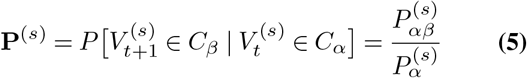

with *α, β* ∈ {1, …, *K*}. For the tentative optimal clustering solution, a TPM was estimated separately for the DTMC of each fMRI scan.

#### Limiting probability

In this study, the limiting distribution was only estimated for irreducible and aperiodic DTMCs (22), with finite state space given by the tentative optimal state trajectories. Therefore, for every subject, *s*, with a DTMC satisfying the aforementioned criteria, it followed that:

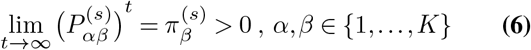

where the estimate of the row vector denoting the stationary distribution of the DTMC (22), 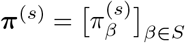, with dimension 1 × |*S*|, is given by:

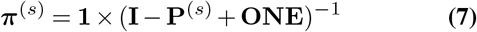

where **1** is a 1 ×|*S*| vector of ones, **I** is the identity matrix with rank |*S*|, **P** ^(*s*)^ is the TPM of subject *s* and **ONE** is a |*S*|× |*S*| matrix all of whose entries are one. Due to the inclusion criteria imposed on the DTMCs defined by the optimal state trajectories, i.e., irreducibility and aperiodicity, only 37 and 46 DTMCs from the HC and SZ groups, respectively, were analyzed. For a given FC state *β, π*_*β*_ (element *β* of the row vector ***π***) was the measure to be used to perform intergroup comparisons. Importantly, since only aperiodic DTMCs were considered, *π*_*β*_ can be understood as the limiting probability that the DTMC is in FC state *β* and as the long-run fraction of time the DTMC spends in FC state *β*. It must be noted that intergroup comparisons between the estimated stationary distributions were not performed in this study.

### Intergroup comparisons

In this research, hypothesis tests to compare the group mean of the properties calculated from the temporal state trajectories observed in SZ patients and HCs were performed using Monte Carlo permutation tests (23) by adapting the procedure used by (12, 17, 18). To produce an accurate approximate estimation of the permutation distribution, these tests were conducted using *B* = 10000 permutations (24). Here, depending on the result of a Levene’s test (used to assess the homogeneity between the group variances) the Monte Carlo permutation tests were performed based on one of the following two statistics (under the null hypothesis):

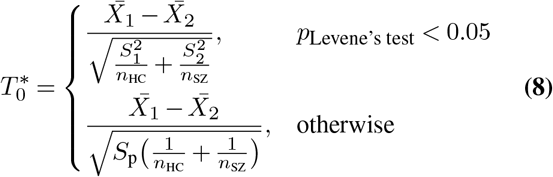

where 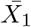 and 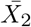 are the random sample means, *S*_1_ and *S*_2_ are the random sample standard deviations and *n*_HC_ and *n*_SZ_ are the sample sizes for the HC and SZ groups, respectively. The pooled random standard deviation, *S*_p_, is given by 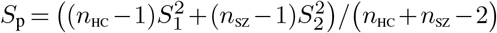. Under the null hypothesis, the statistic from Eq. (8) used to perform the statistical test was subsequently used to obtain the value of the statistic under each of the *B* permutations of the sample data. In this study, the standard deviation of the difference of the group means was estimated using 500 bootstrap samples within each permutation sample. This was performed so that the estimation of this quantity was conducted independently of the calculated means difference.

### Comparison to resting-state functional networks

The functional relevance of the estimated FC states was investigated by assessing whether there was a significant spatial overlap between the centroids/medoids and any of the seven reference RSNs defined by (25). This was accomplished by employing the procedure used by (18, 19). Specifically, the seven RSNs were transformed into seven non-overlapping vectors with dimension 1 × 90, where each entry of the vectors corresponded to the proportion of voxels of each AAL brain area that were assigned to each of the seven RSNs. Finally, the Pearson correlation coefficient was used to assess the spatial overlap between these seven RSNs and the centroids/medoids 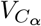 with *α* ∈ {1, …, *K*} (all negative values of 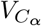 were set to zero so that only areas thought to define relevant functional networks were considered).

### Unsupervised internal cluster validation criteria

The quality of clustering solutions outputted by the clustering algorithms was evaluated using the average Silhouette coefficient and the Dunn’s index (21).

### External validation clustering agreement measures

Clustering outputs from distinct algorithms were compared using the Adjusted Rand Index (ARI) and the Variation of Information (VI) clustering agreement measures (21).

### Clustering stability evaluated by K-fold cross-validation

The stability of clustering solutions was assessed according to a 10-fold cross-validation procedure adapted from (26). Firstly, the sample of the pooled leading eigenvectors was split into two subsamples, referred to as training and test samples. Secondly, a clustering algorithm was applied to the training sample - yielding partition *P*_1_. Subsequently, a Nearest Centroid classifier assigned each observation of the test sample to the cluster of partition *P*_1_, whose centroid was nearest - resulting in the class set *P*_2_ of the test sample. The same clustering algorithm was then applied to the test sample - producing the cluster set *P*_3_. Finally, partitions *P*_2_ and *P*_3_ were compared based on the ARI, VI and percent agreement (fraction of objects correctly assigned). This procedure was repeated for each of the 10 cross-validation folds.

### Software

This analysis used MATLAB R2019b (27), the Statistics and Machine Learning Toolbox™ and the Econometrics Toolbox™.

## Results

### Intergroup differences across partition models detected by the K-means algorithm

The collection of clustering solutions was investigated to search for FC states whose fractional occupancy and dwell time most significantly and consistently differed between SZ patients and HCs. For a partition model with *K* clusters, *K* hypothesis tests were performed. Consequently, to account for the increased probability of false positives, the significance threshold was adjusted to *α*_2_ = 0.05*/K* using a Bonferroni correction. Additionally, a conservative significance threshold of 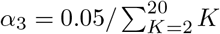 was considered to encompass both dependent and independent null hypotheses across clustering solutions.

Figure 2B presents, for each clustering solution, the *K* two-sided *p*-values obtained from evaluating whether the group mean fractional occupancy of a FC state differed between SZ patients and HCs. From the inspection of Figure 2B, it is apparent that, across all partition models, the clustering procedure consistently returned FC states whose mean fractional occupancy differs significantly between groups - falling below the corrected significance thresholds *α*_2_ and *α*_3_.

**Figure 2.**
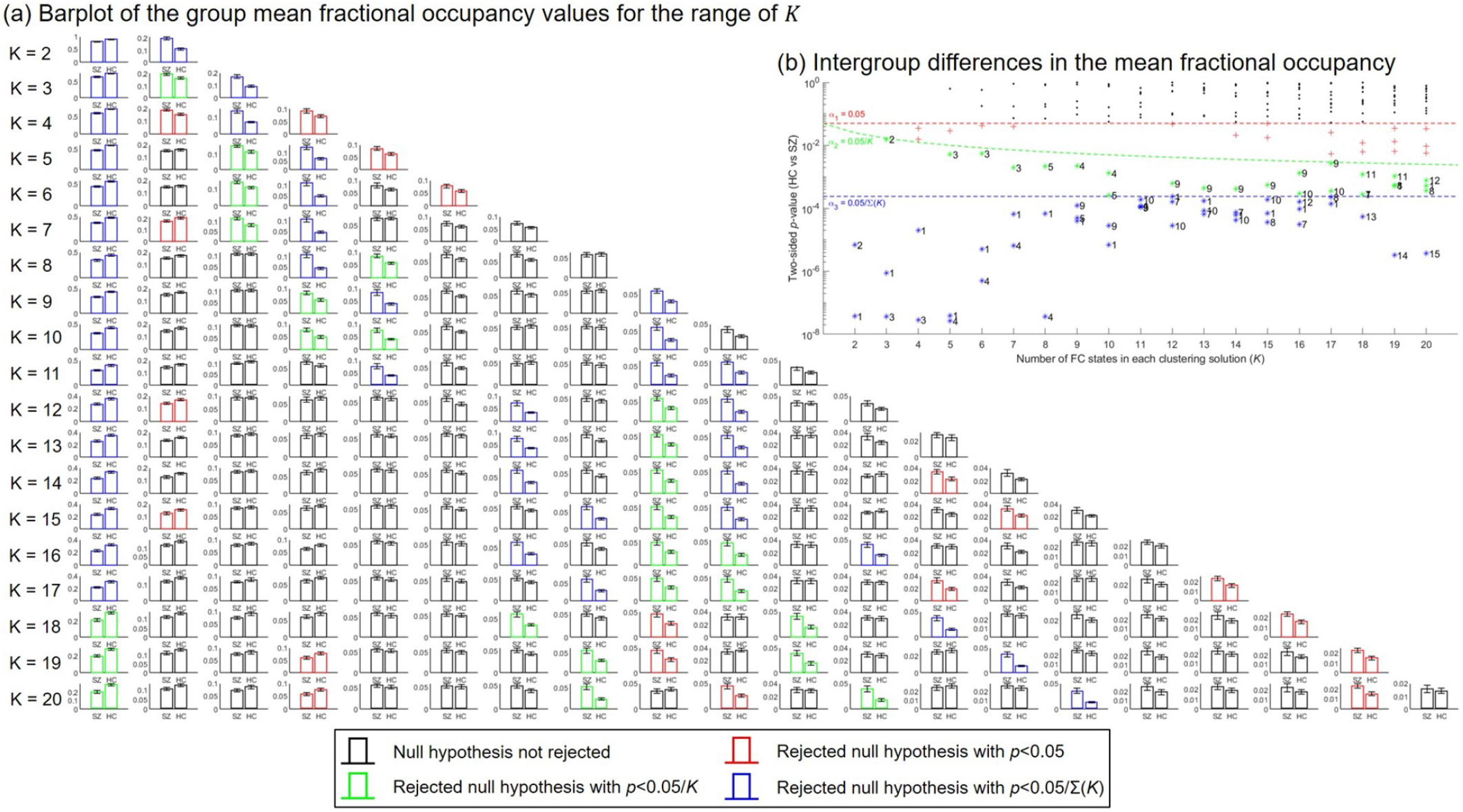
Intergroup comparisons of the mean fractional occupancy of each FC state for each clustering solution. **(A)** Barplot of the estimated mean fractional occupancy with associated standard error of each FC state for each group. For each FC state, the colour of the bars indicates whether the null hypothesis of no intergroup differences in the mean fractional occupancy was rejected (two-tailed tests). The standard error of each bar was calculated as the standard deviation of the sample data divided by the square root of the sample size. **(B)** Two-sided *p*-values obtained for the intergroup comparisons of the mean fractional occupancy of each FC state for each partition model. FC states (clusters) are ranked according to their estimated probability of occurrence, where cluster 1 consists of the largest number of objects and cluster *K* consists of the least number of objects.

Closer inspection of Figure 2B shows there are significant intergroup differences in the mean fractional occupancy of FC state 1 for a range of clustering solutions (*p < α*_3_, two-tailed tests). In fact, the mean fractional occupancy of this state was found to be significantly decreased in SZ patients compared to HCs (*p < α*_3_ for *K* ∈ {2, …, 18}, one-tailed tests), as suggested in Figure 2A. Interestingly, for all partition models, the centroid associated with FC state 1 revealed this recurrent FC pattern represents a global mode of BOLD phase coherence (all elements of the centroid had the same sign). Hence, FC state 1 is referred to as the Global Mode.

As depicted in Figure 2B, across all clustering solutions, further non-global FC states are characterized by significant intergroup differences in the group mean fractional occupancy (*p < α*_3_, two-tailed tests). Interestingly, all these states were typified by a higher mean probability of occurrence in the SZ group compared to the HC group (*p < α*_3_, one-tailed tests), as presented in Figure 2A. Visual inspection of these non-global FC states revealed they represent varying forms of the same underlying connectivity patterns. Specifically, states detected for lower values of *K* could be obtained by combining the fine-grained FC patterns identified in partition models with larger values of *K* - evidencing the dependence among the hypothesis tests performed across clustering solutions.

The analysis of mean dwell time estimates of detected FC states suggested that this measure did not allow as much consistent and clear differentiation between groups compared to the estimates of the fractional occupancy of FC states, as observed in Supplementary Figure S1. In fact, the mean dwell time of the Global Mode was reduced significantly in SZ patients compared to HCs in only 7 clustering solutions (*p < α*_3_, one-tailed tests). Conversely, across all partition models, the mean dwell time was identified as significantly increased in the SZ group compared to the HC group in only two FC states (*p < α*_3_, one-tailed tests). Notably, these non-global FC states were highly correlated (Pearson’s *r* = 0.996) - reinforcing the fact that significant intergroup differences were consistently detected across similar FC patterns.

### Overlap with reference functional networks

Investigation of the overlap between the centroids of the detected FC states and the seven canonical functional networks defined by (25) confirmed that intergroup differences were consistently detected in a number of varying forms of the same FC patterns. Interestingly, FC state 1 did not significantly overlap with any of the seven reference RSNs, as depicted in Supplementary Figure S2A - indicating this global state does not reveal the activation of any particular subset of functionally coupled brain regions. Additionally, across partition models, the non-global FC states with a significantly increased mean fractional occupancy (and dwell time) in SZ were found to repeatedly overlap with the Somatomotor, Dorsal Attention and Limbic networks, as illustrated in Supplementary Figure S2A. Accordingly, FC states with functional activity possibly related to that of the aforementioned canonical RSNs recur more often (and lasted for larger consecutive periods of time) in SZ patients.

### Internal validation of K-means clustering solutions

As shown in Figure 3, the highest average Silhouette coefficient and Dunn’s index were obtained for clustering solutions with a low number of FC states, which are of limited interest for the present study. Contrarily, for clustering solutions with more than 12 clusters, both validation measures remained relatively constant at low values - indicating such partitions are also of limited interest.

**Figure 3.**
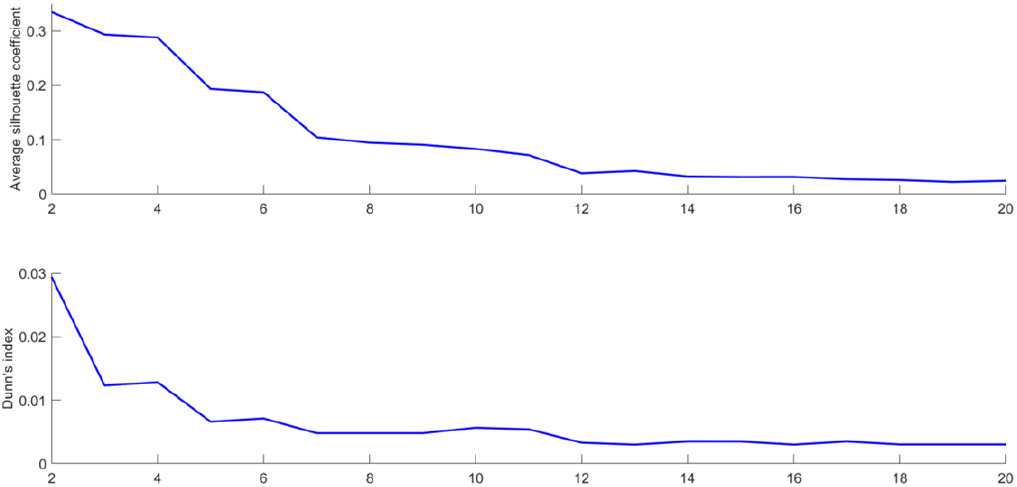
Internal validation of *K*-means clustering results. Average Silhouette coefficient and Dunn’s index used to evaluate the quality of clustering solutions.

Notably, for clustering solutions with *K* between 7 and 11, the average Silhouette coefficient decreased smoothly and the Dunn’s index remained approximately constant, as observed in Figure 3 - suggesting these partition models are of potential interest for further analysis.

### Selection of the optimal clustering solution

For the subsequent analysis, the partition model with 11 FC states was selected as the optimal *K*-means clustering solution. This decision was based upon the ability to identify a collection of FC states with properties that significantly differed between groups and the quality and stability of the actual partition of the data.

The collection of 11 BOLD phase coherence patterns and their fractional occupancy values are presented in Figure 4. The non-global FC states were found to significantly correlate with six of the seven RSNs estimated by (25), as shown in Figure 4A. From Figure 4B, it is apparent that the 11 FC states represent BOLD phase coherence between distinct subsets of brain areas. Furthermore, significant intergroup differences were identified in the mean fractional occupancy of 4 FC states, as observed in Figure 4C. Closer inspection of Figures 4A and 4B reveals that these 4 FC states represent distinct functionally meaningful networks. The mean fractional occupancy of FC state 1 was significantly decreased in SZ patients compared to HCs (Hedge’s *g* = 0.694, medium to large effect size), with estimates 29.8 ± 19.8% and 39±.8 14.9% (mean ± std) for the SZ and HC groups respectively. Furthermore, the mean fractional occupancy of FC states 5, 9 and 10 was significantly increased in SZ patients compared to HCs (Hedge’s *g* = {0.611, 0.630, 0.629}, respectively, medium to large effect size), with estimates 7.61 ± 7.78%, 5.67 ± 6.34% and 5.14 ± 4.09% for the SZ group and 4.02 ± 3.15%, 2.52 ± 3.24% and 2.94 ± 2.81% for the HC group, respectively (mean ±std).

**Figure 4.**
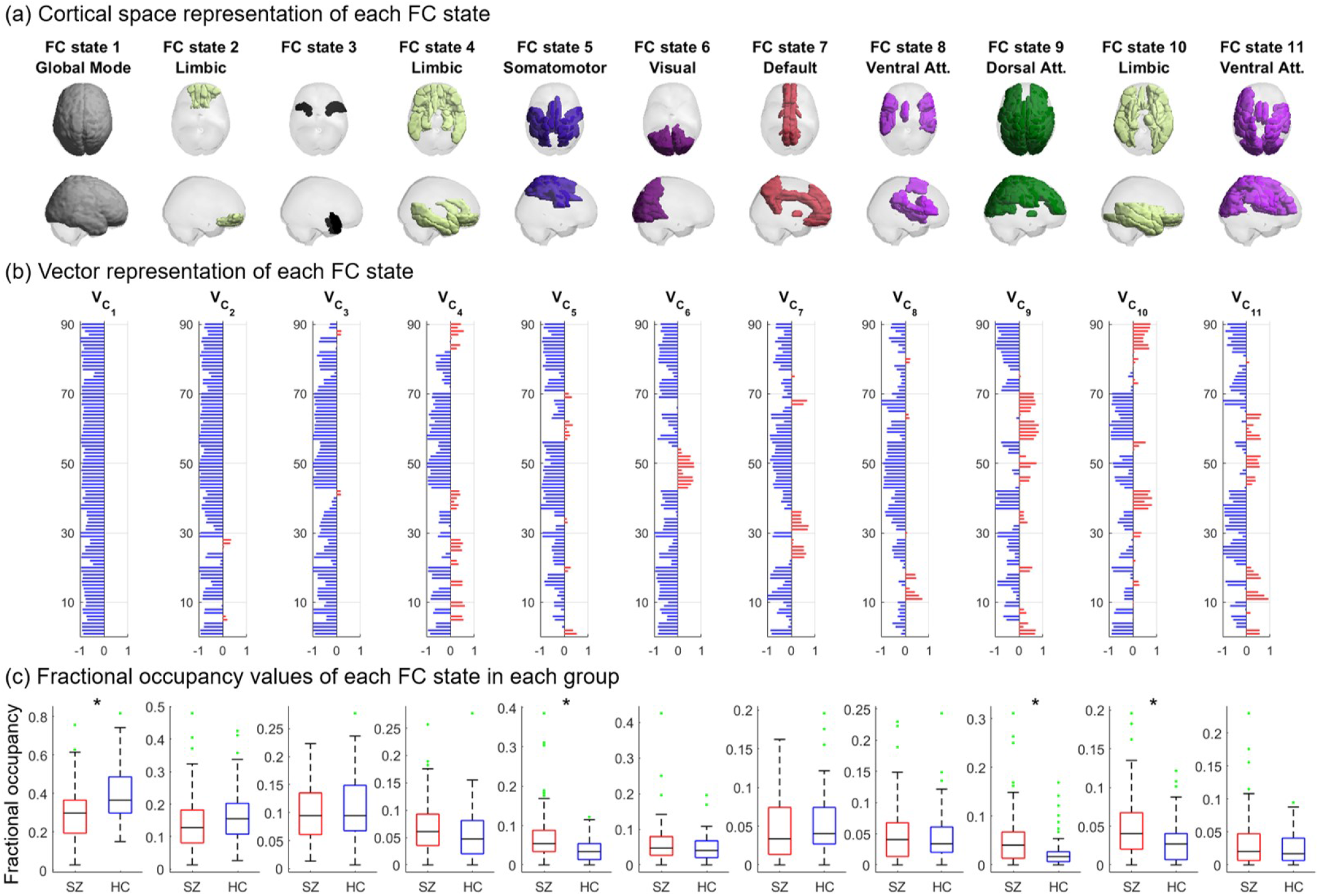
Repertoire of BOLD phase coherence states obtained from the optimal clustering solution with *K* = 11 clusters. The FC states are arranged (left-to-right) according to decreasing estimated fractional occupancy. Each FC state is represented by a *N* × 1 centroid 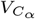, with *α* ∈ {1,…, 11}. **(A)** Cortical rendering of all brain areas with positive values in 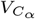. The functional network defined by (25) with which 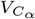 most significantly overlapped is indicated as subtitle. **(B)** Vector representation showing the *N* elements in 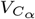, representing the contribution of each brain area to FC state *α*. **(C)** Boxplot of the fractional occupancy values for each FC state for the SZ and HC groups. Asterisks indicate significant intergroup differences (*p < α*_3_, one-tailed tests). Green points represent outliers, according to the Tukey criterion.

### Stability of the optimal clustering solution

The percent agreement, ARI and VI obtained for each fold of the 10-fold cross-validation procedure are provided in Supplementary Table S1. The results suggest respectively, good levels of association and paired agreement between partitions of the test sample and that the amount of information that was lost in changing from the class set *P*_2_ to the cluster set *P*_3_ of the test sample was relatively low. Consequently, the optimal clustering solution is considered valid and appropriate for further analysis.

### State-to-state transitions of the optimal state trajectories

With respect to the optimal clustering solution, for all participants, the individual DTMC defined by the temporal trajectories through the finite state space *S*^′^ = {1, …, 11}, was characterized by its estimated TPM. For each probability of transitioning from state *α* to state *β* (*α*→ *β, α, β* ∈*S*^′^), a two-sided *p*-value was obtained by testing whether its group mean differed between groups. The state-to-state transitions probabilities that were significantly affected in SZ patients compared to HCs are depicted in Figure 5.

**Figure 5.**
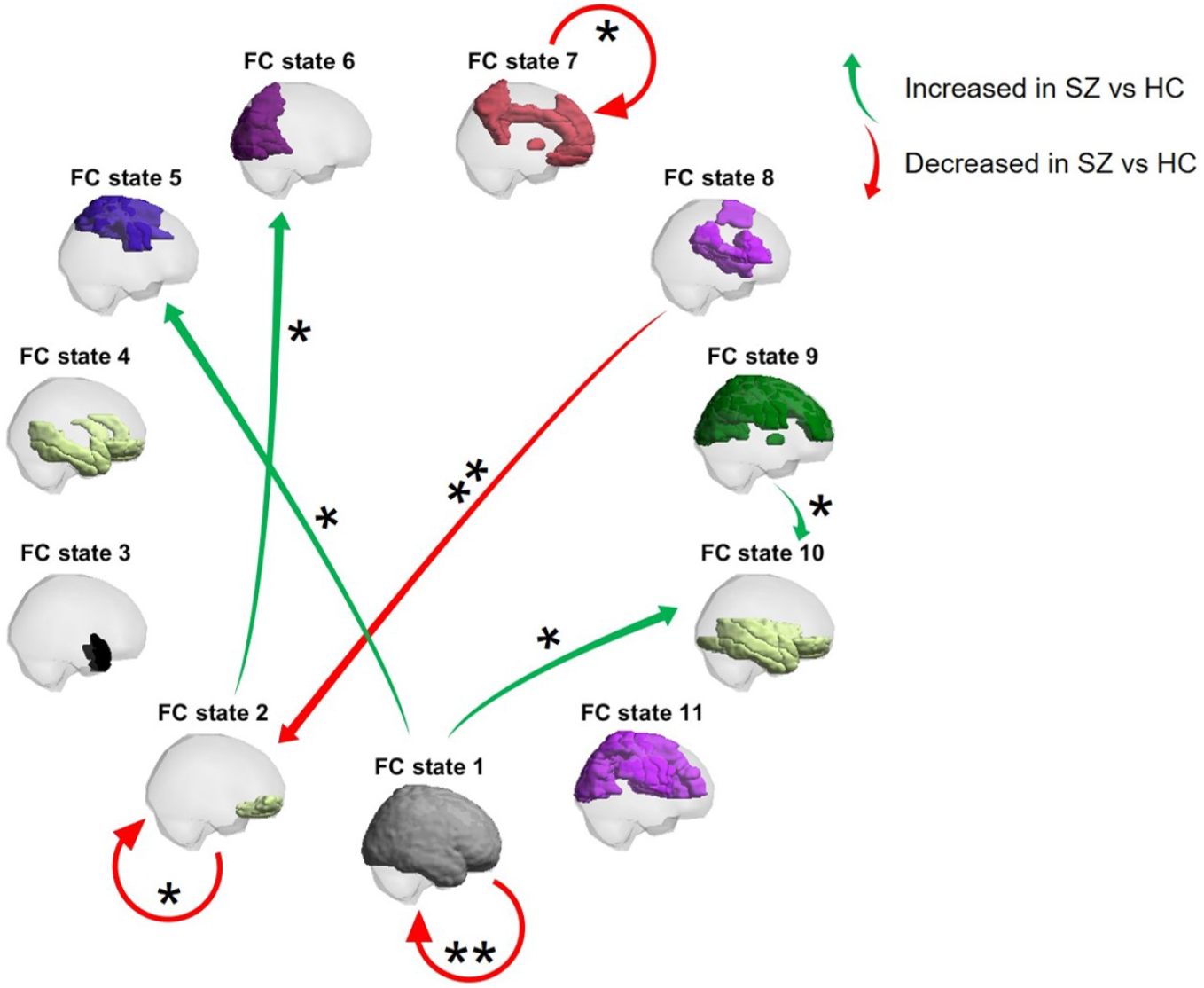
Transition diagram of the state-to-state transitions significantly altered in SZ patients compared to HCs. Arrows represent a mean transition probability that was significantly increased (green) or decreased (red) in SZ patients compared to HCs. Single and double asterisks indicate, respectively, significant intergroup differences with *p <* 0.05*/*11 and *p <* 0.05*/*(11 × 11) (one-tailed tests).

As shown in Figure 5, the mean probability of remaining in FC state 1 was significantly reduced in SZ patients compared to HCs (Hedge’s *g* = 0.726, medium to large effect size). Furthermore, the mean probability of remaining in FC state 7 was significantly reduced in the SZ group compared to the HC group (Hedge’s *g* = 0.515, medium effect size). Lastly, the mean probability of transitioning from FC state 1 to FC states 5 and 10 (Hedge’s *g* = 0.452, 0.513, respectively, small to medium effect size) and from FC state 9 to FC state 10 (Hedge’s *g* = 0.461, small to medium effect size) was significantly increased in SZ patients compared to HCs. Overall, a number of mean transition probabilities were found to be altered in SZ patients.

### Limiting probabilities of the optimal FC states

For the subgroup of 37 HCs and 46 SZ patients with irreducible and aperiodic DTMCs, the estimated mean long-run proportion of TRs spent in FC state 1 was, respectively, 0.309 ± 0.104 and 0.272 ± 0.111 (mean ± std). Surprisingly, no intergroup differences were found in the mean limiting probabil-ity of this state (two-tailed test; Hedge’s *g* = 0.342, small to medium effect size). Only the mean limiting probability of FC states 5 and 10 was identified as significantly increased in the SZ subgroup compared to the HC subgroup (*p <* 0.05, one-tailed tests; Hedge’s *g* = {0.464, 0.449}, respectively, small to medium effect size).

### Influence of using the K-medoids algorithm instead of the K-means algorithm

The application of the *K*-medoids algorithm was found to enable the detection of FC states with a mean fractional occupancy and a mean dwell time that consistently and significantly differ between groups. Similarly with the findings produced by the *K*-means algorithm, the *K*-medoids algorithm identified a FC state which represents a globally BOLD synchronized pattern whose mean fractional occupancy was significantly decreased in SZ patients compared to HCs. Additionally, the mean fractional occupancy of a number of non-global FC states related to the reference Somatomotor, Dorsal Attention and Limbic RSNs was found to be significantly increased in SZ patients compared to HCs.

The ARI and VI showed that the clustering solutions with the same number of FC states detected by each of the clustering algorithms were dissimilar. Interestingly, for each *K*, with *K* ∈ {2, …, 20}, the FC states (centroids/medoids) detected by each of the clustering algorithms with significant intergroup differences in the mean fractional occupancy and mean dwell time (*p < α*_2_, two-tailed tests) were found to be highly correlated. Therefore, both the *K*-means and the *K*-medoids algorithms were found to effectively identify similar FC states whose properties provide the capacity to differentiate SZ patients from HCs.

## Discussion

This study investigated differences between the resting-state dFC observed in SZ patients and HCs.

Across partition models, a globally BOLD synchronized state recurs less in SZ patients. This finding is in line with those of previous studies using different approaches to investigate dFC (6, 7, 9). Conversely, a number of non-global FC states recur more often in SZ patients, as shown in (6, 7, 10). These non-global states, which have been previously referred to as “ghost” attractor states (1, 19), were found to have functional relevance related to that of the Somatomotor, Dorsal Attention and Limbic reference RSNs defined by (25). Notably, despite the lack of a full understanding of the relationship between connectivity patterns observed in Electroencephalography (EEG) and fMRI, these findings could be speculated to portray a temporal dynamics related to that observed with EEG microstates measured at a different time resolution. In fact, in line with the aforementioned findings, EEG studies have reported an increased occurrence of a microstate associated with the Limbic RSN in SZ patients compared to HCs (28). However, the same study showed that the occurrence of a microstate associated with the Attention RSN was decreased in SZ patients (28). This differed from the findings presented here.

From the analysis of the optimal clustering solution, it was found that when in the Global Mode, SZ patients are more likely not to remain in that state in the next time instant, in line with findings from previous studies (6). Importantly, this more frequently occurring pattern of global BOLD phase coherence has been linked to greater neural flexible switching via integration or segregation of different functional connections (12, 29). Therefore, the reduced ability of SZ patients to access and remain in this state could be hypothesized to provoke reduced brain flexibility, i.e., the dynamical state trajectories of SZ patients will, to a greater extent, stay restricted to a fixed set of “ghost” attractor states rather than flexibly transitioning through the full collection of FC states enabled by the recurrent transitions into the functionally integrative Global Mode. This hypothesis seems to be consistent with previous research which found whole-brain integration of higher-order networks was impaired in SZ (8). Lastly, SZ patients were found to transition more likely than HCs from the Global Mode to FC states 5 and 10. It can therefore be suggested that the decreased occurrence of FC state 1 resulted from the increased propensity of SZ patients to transition to networks related to the Somatomotor (state 5) and Limbic (state 10) RSNs. This could have in turn led to their increased occurrence (and limiting probability) - supporting the hypothesis of reduced brain integration of segregated functional connections in SZ.

One interesting finding is that, compared to HCs, SZ patients switch more frequently from a FC state related to the Dorsal Attention RSN (state 9) to a FC state related to the Limbic RSN (state 10). This finding is consistent with that of EEG studies which reported unexpectedly more transitions from a microstate associated with the Attention RSN to a microstate associated with the Limbic RSN (30). Furthermore, SZ patients present a decreased ability to remain in a FC state functionally related to the Default RSN. Notably, this RSN has been linked to core processes of human cognition (2). Therefore, this observation may support the view of SZ as a disorder affecting cognitive function. Additional intergroup differences were detected in a number of other state-to-state transition probabilities. It may be the case therefore that the observed abnormal dynamical state transitions provide potential biomarkers of this disorder.

On the question of the influence of using the *K*-medoids algorithm to conduct a LEiDA analysis, this study found that similar intergroup differences are captured by employing either the *K*-medoids algorithm or the *K*-means algorithm. This finding suggests that the choice of an optimal clustering algorithm should rely not only on statistical and cluster validation analyses, but also on concepts and methods from dynamical systems theory (1, 12, 19). On the one hand, from the definition of the *K*-means algorithm, the detected FC states (centroids) are not necessarily observations from the input dataset, but could rather be interpreted as averaged recurrent unobserved FC patterns; hence their designation as “ghost” attractor states (1, 19). However, the definition of the *K*-medoids algorithm implied the detected FC states (medoids) are observed recurrent FC patterns. Research questions pertaining to the functional meaning of the detected FC patterns underline the need to employ tools from dynamical systems theory to provide further insights into the dynamical regime of brain activity (1, 12, 19).

Developing on from previous LEiDA analyses, this study proposes examining the limiting probability of FC states. This property offers valuable insights into the long-run proportion of time that a DTMC spends in each state. Specifically, this measure is computed from the TPMs which characterize the state trajectories - capturing dynamic behaviour of brain activity to a greater extent than fractional occupancy. However, considerably more research will need to be conducted to determine its utility. Furthermore, the measurements of this property are derived from the estimation of the stationary distribution of the state trajectories, defined as irreducible and aperiodic DTMCs. A natural progression of this work is to examine whether intergroup differences in the stationary distributions provide further insights into the limiting dynamic behaviour of brain activity in diseased and healthy populations. This could be achieved by employing the two-sample goodness of fit *χ*^2^ test.

One shortcoming of this study which could have affected the measurements of both the state-to-state transition and state limiting probabilities is the low temporal resolution of the neuroimaging data (TR = 2 s). This was most clearly observed from the inconsistencies found across state trajectories obtained from the optimal clustering solution where, often times, the occurrence of all FC states was not guaranteed. In fact, Magnetoencephalography (MEG) studies have suggested that brain functional connectivity dynamics occurs at time scales of approximately 200 ms (31, 32). Accordingly, future work should utilize data with higher temporal resolution to enable the capture of more rapid dynamics - improving the utility of these properties as possible biomarkers of SZ.

Another limitation of this study is that the detected FC states were strongly constrained by the selected parcellation atlas (AAL). Despite having shown consistent results across studies employing LEiDA (12, 17–19), the AAL template is based on an anatomical parcellation and, therefore, may not provide an adequate framework to conduct an analysis of dFC. Accordingly, future studies could extend this analysis to other fMRI-derived anatomical or functional parcellations.

An important limitation lies in the fact that the effect of variables such as age, gender and clinical history of patients was not taken into account while assessing intergroup differences - hindering the identification of reliable biomarkers of SZ (33). Specifically, intergroup differences were attributed only to the effect of the group. Further research is required to determine whether these variables or their interaction could explain the variability found between groups. Another issue that was not addressed in this work was whether not applying temporal filtering and nuisance regression strategies influenced the LEiDA method and therefore, the observed intergroup differences. Further research on this question could contribute with valuable insights into these controversial preprocessing steps.

Finally, this study provides unbiased and statistically rigorous evidence for differences between SZ patients and HCs. Nevertheless, the potential to serve as biomarkers of SZ and the clinical implications of the results derived herein remain to be analyzed. Future investigations should gather a diverse panel of experts to explore how these findings could be applied to improve our understanding of SZ.

## Conclusions

Resting-state dynamic functional connectivity comparisons were conducted between SZ and HCs by employing and extending the LEiDA method. Through the characterization of the temporal expression of different FC patterns, this study found that SZ patients exhibit a reduced capacity to access and remain in a globally BOLD synchronized state. An implication of this is the possibility that, even in the absence of any explicit task, SZ patients transition more frequently to network patterns that are commonly activated during specific tasks.

## Author contributions

MF carried out the analysis and wrote the main manuscript. JC and CA verified and advised the theoretical methods. JC and CA supervised the whole project. All authors participated in the discussion of the ideas. MF and JC contributed in the final writing of the manuscript.

## Acknowledgments

The imaging data was collected and shared by the Mind Research Network and the University of New Mexico funded by a National Institute of Health Center of Biomedical Research Excellence (COBRE) grant 1P20RR021938-01A2. This work has been funded by National funds, through the Portuguese Foundation for Science and Technology (FCT) - projects UIDB/50026/2020 and UIDP/50026/2020. JC is funded by FCT grant CEECIND/03325/2017.

## Supplementary Note 1: Analysis of dwell time estimates of FC states detected by using the K-means algorithm

**Figure S1.**
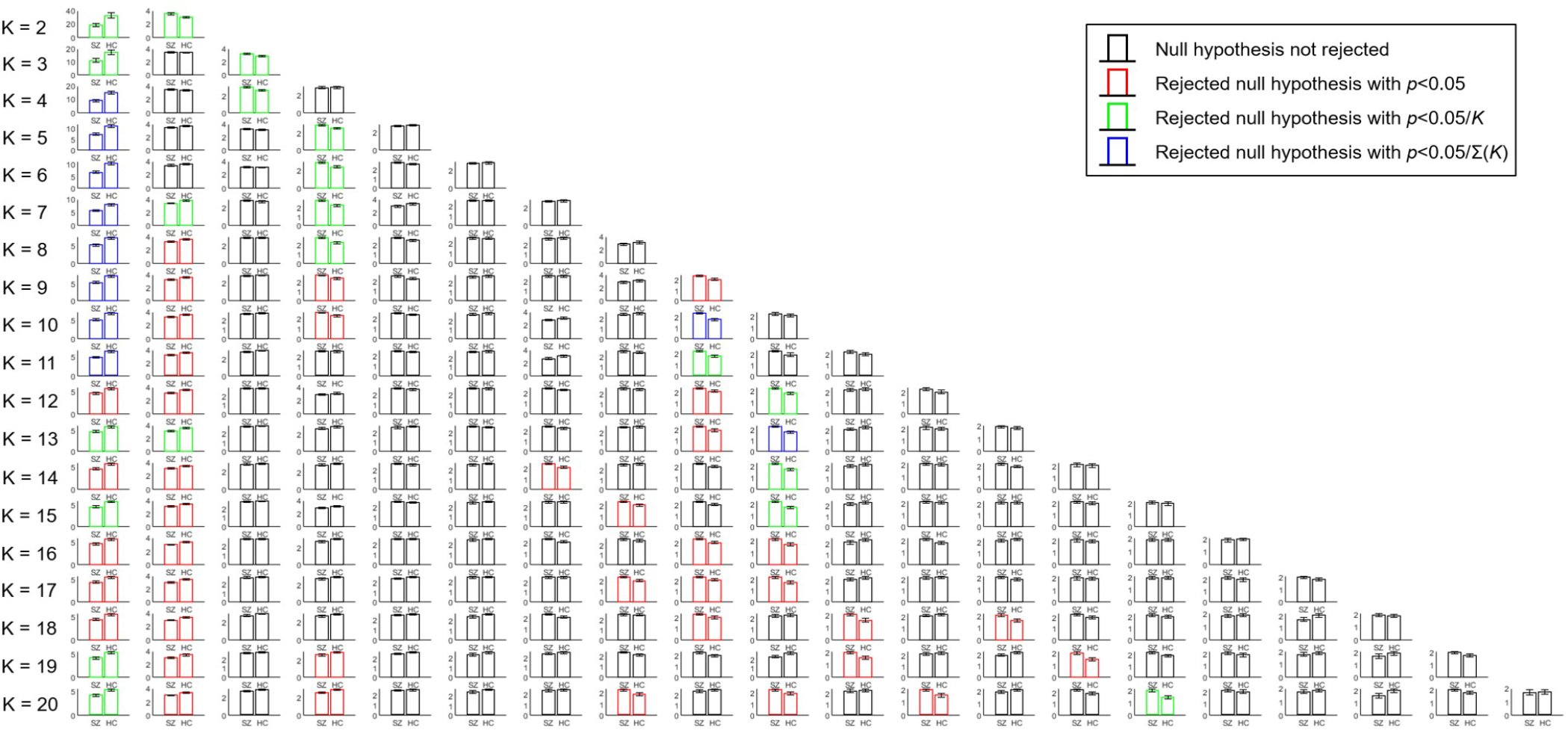
Intergroup comparisons of the mean dwell time of each FC state for each clustering solution. Barplot of the estimated mean dwell time with associated standard error of each FC state detected by the *K*-means algorithm for each group. For each FC state, the colour of the bars indicates whether the null hypothesis of no intergroup differences in the mean dwell time was rejected (two-tailed tests). Black bars indicate the null hypothesis was not rejected at a 5% significance level. Red, green and blue bars indicate the null hypothesis was rejected at a 0.05, *α*_2_ and *α*_3_ significance level, respectively. The standard error of each bar was calculated as the standard deviation of the sample data divided by the square root of the sample size.

## Supplementary Note 2: Overlap of cluster centroids with reference resting-state networks

**Figure S2.**
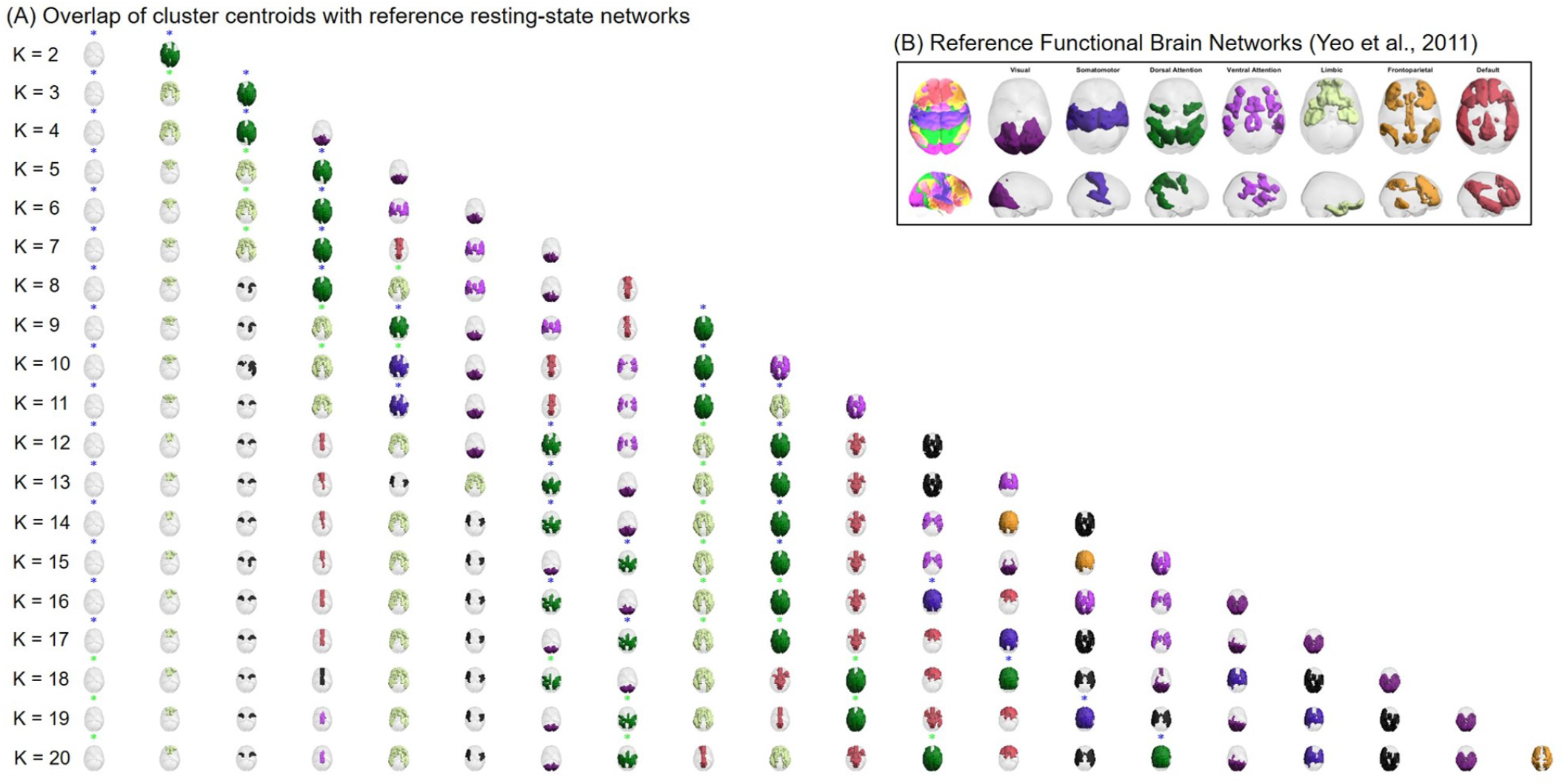
Overlap between FC states and Functional Brain Networks. **(A)** Representation of the centroids obtained for each clustering solution in cortical space. The rendered brain areas correspond to positive elements in the vectors of the centroids. Brain regions are coloured according to the reference RSNs defined by (25) whose *p*-value obtained from computing the Pearson correlation coefficient was lowest (with *p <* 0.05*/K*). Centroids not significantly overlapping with any of the reference RSNs are coloured in black. For each FC state, title asterisks indicate whether significant intergroup differences in the mean fractional occupancy were detected. Green and blue asterisks indicate *p < α*_2_ and *p < α*_3_ (two-tailed tests), respectively. **(B)** Reference functional brain networks estimated by (25).

## Supplementary Note 3: Stability of optimal K-means clustering solution

**Table S1.** Stability analysis of the optimal clustering solution. Results obtained across the 10 cross-validation folds for each clustering agreement measure.

| Indices | Cross-validation fold number |  |  |  |  |  |  |  |  |  |
| --- | --- | --- | --- | --- | --- | --- | --- | --- | --- | --- |
|  | 1 | 2 | 3 | 4 | 5 | 6 | 7 | 8 | 9 | 10 |
| Percent agreement | 0.642 | 0.536 | 0.583 | 0.688 | 0.529 | 0.564 | 0.602 | 0.698 | 0.547 | 0.523 |
| Adjusted Rand | 0.692 | 0.712 | 0.717 | 0.757 | 0.696 | 0.678 | 0.749 | 0.718 | 0.773 | 0.709 |
| Variation of information | 1.379 | 1.504 | 1.349 | 1.237 | 1.396 | 1.402 | 1.291 | 1.229 | 1.181 | 1.388 |

## Notes

### Competing Interest Statement

The authors have declared no competing interest.

